# Control of brain state transitions with light

**DOI:** 10.1101/793927

**Authors:** Almudena Barbero-Castillo, Fabio Riefolo, Carlo Matera, Sara Caldas-Martínez, Pedro Mateos-Aparicio, Julia F. Weinert, Enrique Claro, Maria Victoria Sánchez-Vives, Pau Gorostiza

## Abstract

Behavior is driven by specific neuronal activity and can be directly associated with characteristic brain states. The oscillatory activity of neurons contains information about the mental state of an individual, and the transition between physiological brain states is largely controlled by neuromodulators. Manipulating neural activity, brain rhythms or synchronization is of significant therapeutic interest in several neurological disorders and can be achieved by different means such as transcranial current and magnetic stimulation techniques, and by light through optogenetics, although the clinical translation of the latter is hampered by the need of gene therapy. Here, we directly modulate brain rhythms with light using a novel photoswitchable muscarinic agonist. Synchronous slow wave activity is transformed into a higher frequency pattern in the cerebral cortex both in slices *in vitro* and in anesthetized mice. These results open the way to the study of the neuromodulation and control of spatiotemporal patterns of activity and pharmacology of brain states, their transitions, and their links to cognition and behavior, in different organisms without requiring any genetic manipulation.

## INTRODUCTION

All perceptions, memories, and behaviors are rooted in the communication between the billions of neurons that constitute the brain.^1–3^ Individual neurons transmit information using chemical and electrical signals, and are organized in groups or circuits involved in different functions.^3–5^ The electrochemical interactions of these neuronal ensembles produce synchronized electrical activity in the brain, that constitute brain rhythms or oscillations due to their periodic properties, and brain waves when those oscillations show spatial propagation.^6–8^ Brain rhythms can be recorded by different means, involving largely electrophysiological (e.g. electroencephalography (EEG), local field potential (LFP), etc.) or imaging methods. Brain rhythms, their frequency, synchronization or functional connectivity associated to them, are associated with internal brain states and result in specific behaviors.^1^ Neuronal and network activity undergoes significant changes during brain and behavioral state transitions, which have been linked to substantial changes in EEG pattern activity.^9^ For example, large and synchronous brain waves are mostly associated with deep sleep, whereas in wakefulness there is a shift towards more desynchronized and short-amplitude wave patterns.^7,10,11^ These changes in brain and behavioral states, and the concomitant alterations in EEG activity, can be driven by the action of neuromodulators like acetylcholine (ACh).^9,12^ However, it is not fully known how the different cells expressing ACh receptors contribute to the alteration of the global cortical state. Cholinergic receptors include ionotropic nicotinic ion channels and muscarinic metabotropic G protein-coupled receptors. Together, they modulate cortical activity on a fine spatial scale and are involved in crucial neocortical functions such as attention,^13,14^ learning,^15–17^ memory,^18^ and sensory and motor functions.^19,20^ In the neocortex, ACh is released mostly at cholinergic afferents from neurons distributed within the basal forebrain (BF) nuclei. Electrical stimulation of the nucleus basalis can evoke the release of ACh in the neocortex but in an unselective manner, as ascending projections from BF nuclei not only comprise cholinergic axons, but also GABAergic and glutamatergic axons.^21^ Such a lack of selectivity complicates the study of cholinergic signaling in the neocortex and its effects on controlling brain states.

Selective stimulation of cholinergic projections in the neocortex from BF nuclei has been demonstrated with optogenetics, which enables the disruption of neocortical synchronous activity during certain sleep states. Optogenetic stimulation of BF cholinergic neurons also revealed their influence in awake cortical dynamics, coding properties of V1 neurons, and the importance of cholinergic neuromodulation for visual discrimination tasks, showing that stimulation of BF cholinergic cells activates cortical transitions faster than previously presumed.^10^ However, the cell type specificity of optogenetics is limited by the availability of suitable promoters. In addition, it is based on the overexpression of microbial proteins using genetic manipulation, which can distort synaptic physiology.^22–24^ This raises safety and regulatory concerns regarding therapeutic applications.

The control of neuronal signaling with photopharmacology is based on synthetic ligands that target endogenous proteins, and thus its physiological relevance spans from circuit to sub-cellular levels. Since neuronal receptors are highly conserved, photoswitchable ligands can generally be used in multiple species, and their safety and regulation can be established in the same manner as other drugs. The cholinergic system is key to the modulation of a variety of CNS functions,^25^ and thus the use of selective and photoswitchable cholinergic drugs to spatiotemporally control cortical activity could have relevant scientific and clinical implications. Herein, we report that the cholinergic-dependent brain state transitions in the neocortex can be directly controlled using Phthalimide-Azo-Iper (PAI)^26^ a photoswitchable agonist that targets M2 muscarinic acetylcholine receptors (mAChRs) without requiring electric or genetic manipulation. In particular, PAI enables the modulation of spontaneous emerging slow oscillations (SO) in neuronal circuits. PAI *cis-to-trans* photoisomerization decreases the Down- and Up-state durations and increases oscillatory frequency in cortical slices. In addition, PAI allows the reversible manipulation of the cortical oscillation frequency in anesthetized mice using light. Thus, photopharmacology allows the selective control of slow oscillations *in vitro* and *in vivo*, opening the way to the analysis of their spatiotemporal dynamics and their effects on brain and behavioral state transitions.

## RESULTS

### Non-specific activation of mAChRs evokes neuronal hyperexcitability in cortical slices

Changes in cortical rhythms underlie behavioral state transitions, and endogenous ACh actions play a central role in such variations.^27–33^ However, a complete and unifying view has not yet emerged^11^ regarding the cholinergic impact on neuronal and synaptic physiology, and thus on neocortical network dynamics. It is known that ACh contributes to the shift of the neocortical network state from synchronous to asynchronous activity-associated to awake state-in a dose-dependent manner, but the activity of ACh in the vast majority of neocortical neurons and synapses is still poorly characterized. Neocortical activation of mAChRs *in vitro* facilitates synaptic transmission,^34^ recurrent excitation,^35^ and reversibly increases the power of fast-frequency oscillations.^36–39^ On these basis, we first studied *in vitro* the effect of Iperoxo, a potent muscarinic agonist.^40,41^ The goal was to evaluate the potential of mAChRs to selectively modulate the dynamics of isolated neocortical slices, while avoiding the simultaneous activation of nicotinic cholinergic receptors (nAChRs), and to validate Iperoxo photoswitches and useful photopharmacological tools to control neuronal activity. Isolated cortical slices spontaneously generate cortical SO, a hallmark of activity during deep sleep or anesthesia.^42^ We recorded the spontaneous oscillatory activity (control) and then the activity under different concentrations (1, 10, 100 nM) of Iperoxo. The spontaneous slow oscillatory activity alternates between periods of activity or high neuronal firing (Up states) and periods of near silence (Down states). The activation of mAChRs by Iperoxo resulted in a global change in the network’s dynamics to a hyperexcitable state (Fig. 1). At 100 nM Iperoxo, the oscillatory frequency increased (from 0.89 ± 0.12 Hz in the control, to 1.34 ± 0.18 Hz with 100 nM Iperoxo), and the relative firing rate^43^ of the Up-states decreased (from 1.13 ± 0.23 a.u. to 0.10 ± 0.02 with 100 nM Iperoxo) (Fig. 1B). Furthermore, at concentrations equal or higher than 100 nM Iperoxo, the oscillatory activity evolved to periods of seizure-like discharges (Fig.1C),^44,45^ characterized by low (<1 Hz), delta (1-4 Hz) and alpha-beta (8-16 Hz) frequencies (Fig. 1D).

**Figure 1:**
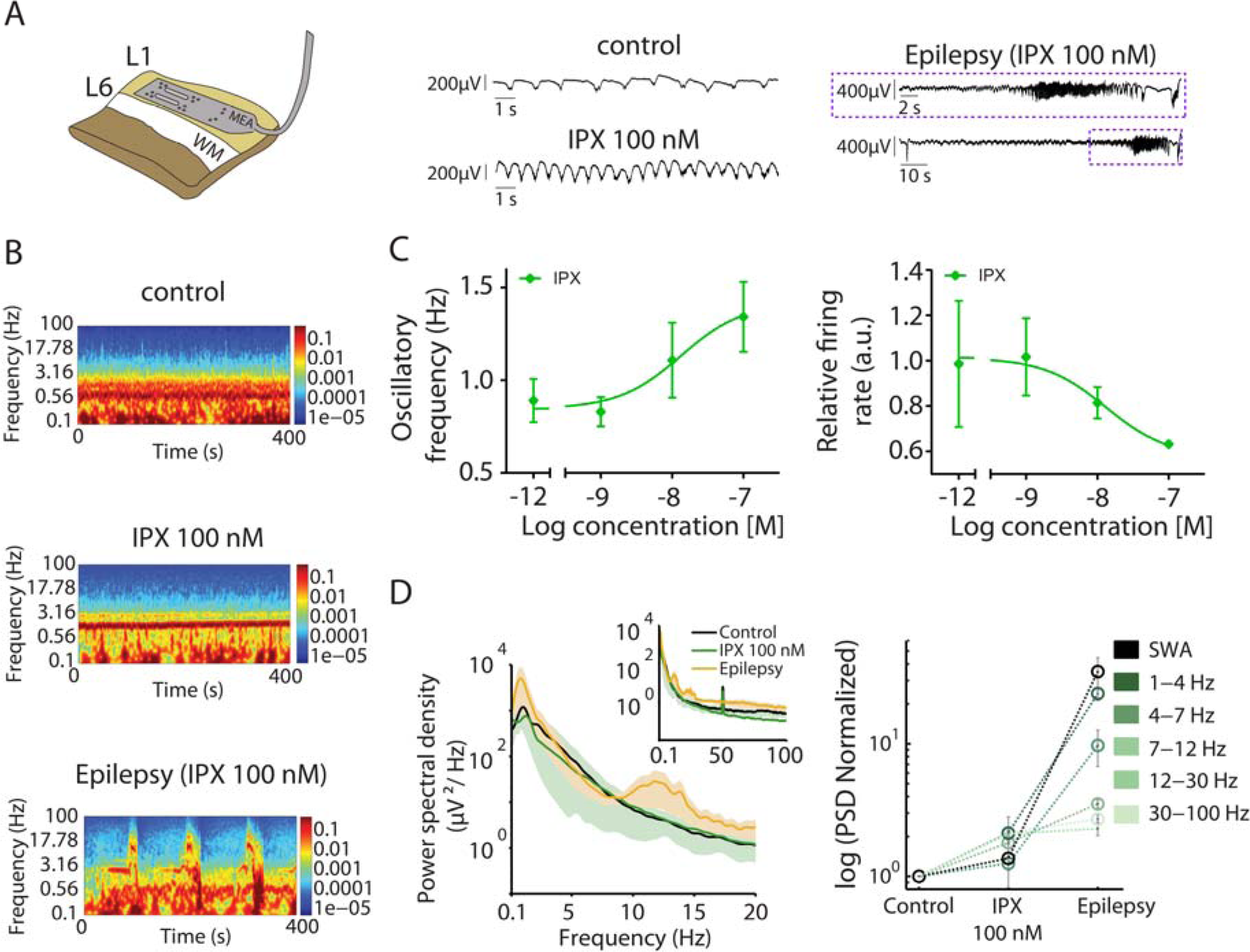
Non-specific activation of mAChRs evokes neuronal hyperexcitability in cortical slices. **(A)** On the left, the experimental setup: MEA, multielectrode array; WM, white matter; L1-L6, layer 1-6. On the right, representative Local Field Potential (LFP) traces showing the increasing of the OF corresponding to the Spectrogram of panel **B. (B)** Spectrogram under color from the same time recording of LFP traces on panel **A**: control, 100 nM Iperoxo and periods seizure-like discharges. (**C**) OF (Hz) and relative firing rate (a.u.). (**D**) Power spectral density (PSD) values showing low, delta and theta frequency component. Normalized PSD values at different frequency bands.

### Effect of PAI isomers on slow and fast oscillations *in vitro*

The hyperexcitable network state induced with Iperoxo reflects the impact of mAChRs activation on cortical networks and brain states.^11^ In order to remotely control these states, we aimed at the observed muscarinic neuromodulation using PAI, a novel photoswitchable Iperoxo derivative that allows the reversible activation of M2 mAChRs with light.^26^ M2 receptors play a relevant role in several CNS disorders,^25^ and regulating their activity and subsequent effects on cortical neuronal networks may provide new therapeutic opportunities for these diseases involving the cholinergic system.

Taking advantage of its profile as a M2 light-regulated agonist, we studied its effect on the modulation of neuronal oscillations in cortical slices. Thus, we applied PAI to spontaneously active neocortical brain slices and recorded their oscillatory activity before and after activating the drug by means of illumination. We first generated the dose-response curves of the two drug forms separately, *trans*-(dark-adapted state) and *cis*-PAI (after UV irradiation), in order to discern the differences in cortical activity between the PAI-isoforms, and to identify the most convenient concentration range to manipulate brain waves with light. The baseline activity (characterized by slow oscillations) was recorded as a control, prior to bath-application of solutions with increasing PAI concentrations (10 nM, 100 nM, 300 nM, and 1 µM, *n* = 6 for each PAI form, *trans* and *cis*) (Fig. 2A). Comparing with the control situation, the *trans*-PAI activity showed alterations in the Up- and Down-state sequence already at 100 nM (Fig.2), leading to an increment in the oscillatory frequency (from 0.58 ± 0.06 Hz during control to 1.87 ± 0.12 Hz applying 1 µM *trans*-PAI) (Fig. 2B). Moreover, increasing concentrations of *trans*-PAI decreased the relative firing rate during the Up states (from 0.98 ± 0.11 a.u. during control to 0.37 ± 0.03 a.u. with 1 µM *trans*-PAI) (Fig. 2C, D). In comparison, *cis*-PAI displayed significantly weaker effects, in agreement with the reported PAI properties.^26^ At 100 nM and 300 nM, *cis*-PAI did not alter the spontaneous activity observed in control experiments (control oscillatory frequency: 0.48 ± 0.037 Hz; 100 nM *cis*-PAI OF: 0.52 ± 0.07 Hz; 300 nM *cis*-PAI OF: 0.87 ± 0.2 Hz), in contrast to the strong alterations in oscillatory activity obtained with 100 nM and 300 nM *trans*-PAI (Fig. 2). Only at concentrations as high as 1 µM, *cis*-PAI altered the Up- and Down-state sequence in comparison to the control, increasing the oscillatory frequency (1.32 ± 0.27Hz) and decreasing the relative firing rate of the Up-states (from 0.86 ± 7 0.04 a.u. during control to 0.74 ± 0.27 a.u. with 1 µM *cis*-PAI) (Fig. 2C, D). In summary, the most compelling differences between *trans*- and *cis*-PAI emerged between 100 nM and 300 nM and were observed in the oscillatory frequency and in the Up-states’ relative firing rate (a.u.) (Fig. 2). We focused on these concentration range therefore in order to photomodulate cortical slow oscillations using PAI.

**Figure 2:**
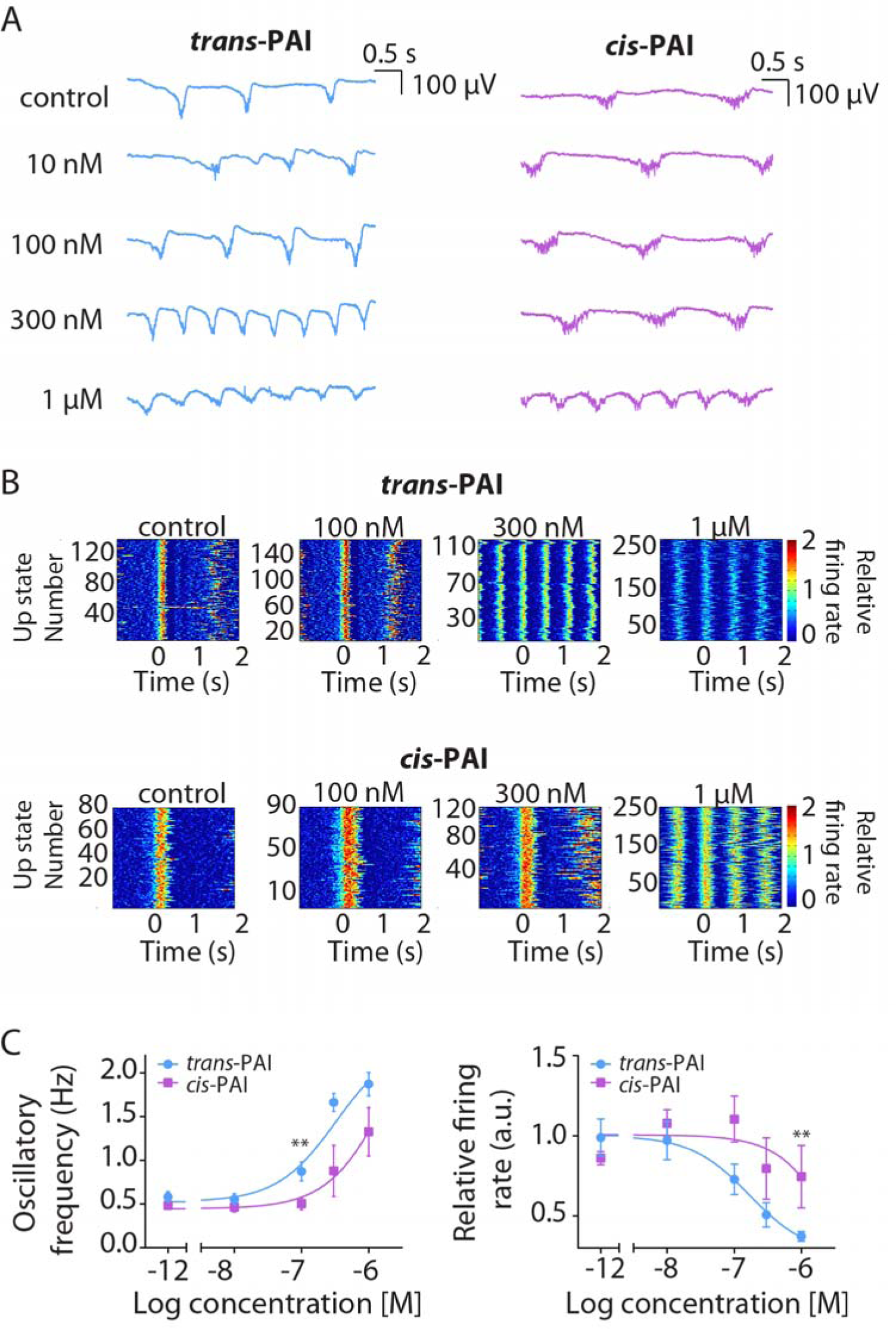
Effect of mAChRs activation by *trans*-PAI and *cis*-PAI on SO. **(A)** Raw LFP example recordings showing the different ability of *trans*- and *cis*-PAI to increasing the OF. **(B)** Raster plots showing the relative firing rate (color coded) under control conditions and different *trans*- and *cis*-PAI concentrations. (C) OF (Hz) and Relative firing rate (a.u.) of the two different PAI isomers, *trans*-(blue, n = 6) and *cis*-PAI (pink, *n*=6) at different concentrations. (\*\**p*<0.01).

### PAI effectively modulates cortical slow oscillations *in vitro*

Once the different oscillatory activity evoked by *cis*- and *trans*-PAI was quantified *in vitro* (Fig.2), we moved on to the control of the rhythmic activity with light in the same cortical slices (Fig.3). We took advantage of the thermal stability of both PAI forms to apply initially the inactive one (*cis*-PAI) at 200 nM in cortical slices (*n* = 15), in the absence of white light to avoid photoconversion to *trans*-PAI during the recordings.^26^ As shown in Fig.2, 200 nM *cis*-PAI evoked an increase in the oscillatory frequency (from 0.53 ± 0.04 Hz in control conditions to 1.04 ± 0.14 Hz with *cis*-PAI, *p*-value = 0.0016), and no significant effects in the relative firing rate of the Up-states (from 0.98 ± 0.09 a.u. in the control to 0.86 ± 0.10 a.u. with *cis*-PAI, *p*-value = 0.2868) (Fig. 3). Subsequent illumination of the slices with white light produced a robust increase in the oscillatory frequency (from 0.53 ± 0.04 Hz in the control to 1.68 ± 0.13 upon illumination, *p*-value = 2.93 * 10^−4^; from 1.04 ± 0.14 Hz with *cis*-PAI to 1.68 ± 0.13 upon illumination, *p*-value = 3.51 * 10^−4^), a significant decrease in relative firing rate of the Up-states (from control values of 0.98 ± 0.09 a.u. to 0.52 ± 0.06 a.u. upon illumination, *p*-value = 2.93 * 10^−4^; from 0.86 ± 0.10 a.u. with *cis*-PAI to 0.52 ± 0.06 a.u. upon illumination, *p*-value = 2.93 * 10^−4^), and a noticiable change in the activity regime of the network (Fig. 3 A and B). The changes in the power spectrum in the population were incremental, as illustrated in the normalized power spectrum (Fig. 3). These changes are similar to the effects of adding Iperoxo to the bath (Fig. 1), and in agreement with PAI photoconversion to the active form (*trans*). The modification in cortical activity was not reversible with 365 nm light (to isomerize PAI to the *cis* form *in situ*), due either to the irreversibility of muscarinic stimulation in the absence of BF afferents and other circuit components in brain slices, or to neuronal potential changes produced by UV light.

**Figure 3:**
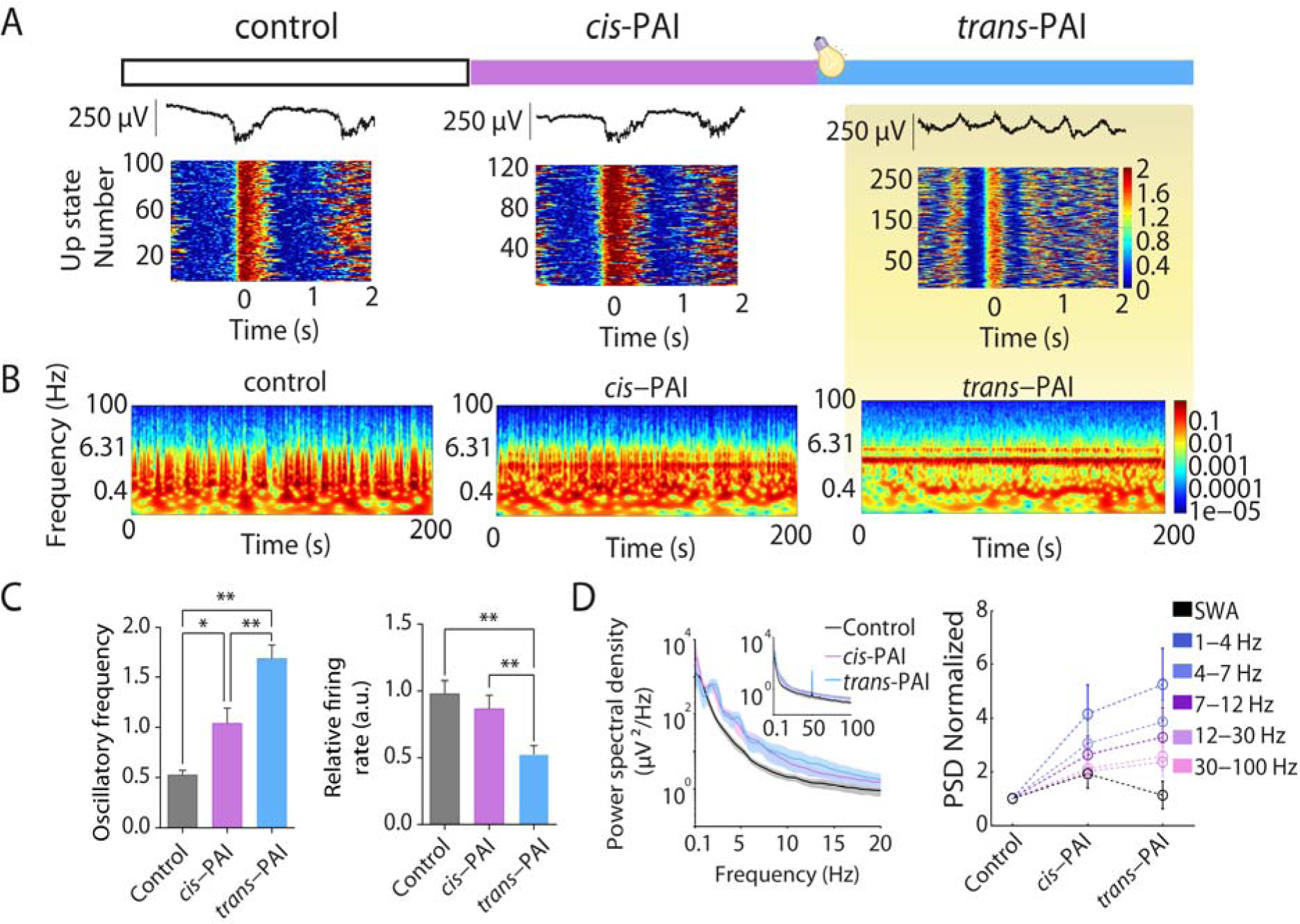
Modulation of brain waves *in vitro* using PAI, a light-regulated ligand. **(A)** Representative Local Field Potential traces (top) and raster plots of relative firing rate under control conditions, 200 nM of *cis*-PAI and 200 nM *trans*-PAI after light activation (n = 15) (bottom). **(B)** Representative spectrogram under control condition, 200 nM of *cis*-PAI and 200 nM *trans*-PAI after light activation. **(C)** OF (Hz) and relative firing rate of the Up-states (a.u.) at 200 nM PAI after pre-irradiation at 365 nm (*cis*-PAI), and photoswitching with white light (*trans*-PAI). **(D)** Averaged Power Spectral Density (PSD) under control conditions, 200 nM of *cis*-PAI and 200 nM *trans*-PAI after white light activation (color code). \**p*<0.005 and \*\**p*<0.001.

### PAI can modulate brain wave activity with light *in vivo*

Having established the unique ability of PAI to alter cortical oscillatory activity with light in slices, we then aimed to photocontrol brain state transitions *in vivo*. Cortical brain waves were recorded in C57BL6/JR mice (*n* = 4) with an electrode inserted through a craniotomy across which we carried out the drug application and brain illumination (see Methods). Initially, we induced deep anesthesia in the animals, a state that is known to reproduce the slow wave sleep state,^28,29,46^ and which is characterized by the generation of cortical slow oscillations similar to the slow frequency waves observed in our experiments in slices under control conditions (Fig.1–3).^6^ Such slow oscillations activity in anesthetized mice was recorded for 500 s under white light illumination of the brain, and the characteristic parameters obtained (oscillatory frequency 0.64 ± 0.06 Hz, relative firing rate 0.83 ± 0.23 a.u.) were taken as the control condition *in vivo*. As the dose-response curves of *cis*- and *trans*-PAI could be different from the *in vitro* conditions (Fig. 2), we tested two different concentrations, 200 nM and 1 µM. 100 µL of 200 nM *cis*-PAI solution were initially applied to the brain surface, and the activity was recorded for another 500 s in the absence of white light, to avoid *cis* to *trans* photoisomerization of PAI. The oscillatory frequency and the frequencies power of the cortical oscillatory activity was not significantly altered by *cis*-PAI (0.61 ± 0.06 Hz) and caused a minor increase in the relative firing rate (1.02 ± 0.31 a.u.). Subsequently, we illuminated the brain using white light in the proximity of the recording electrode, in order to isomerize PAI to its active form (*trans*) and recorded the activity for another 500 s. An increase in the oscillatory frequency to 0.76 ± 0.1 Hz and marked power in the 1 Hz band was observed under white light illumination, without changes in the relative firing rate (1.04 ± 0.27 a.u.). Applying a higher concentration of *cis*-PAI (1 µM) slightly reduced the OF to 0.61 ± 0.1, decreased power at 1Hz band without significantly affecting the relative firing rate (0.99 ± 0.27 a.u.). However, following illumination, the oscillatory frequency increased to 0.83 ± 0.06 Hz, appeared again 1 Hz frequency band and the relative firing rate decreased to 0.86 ± 0.19 a.u. (Fig. 4), following the tendency previously observed *in vitro*. By using two concentrations of the drug, we observed that the effect on the oscillatory frequency was already reached with 200 µM, while the tendency of the firing rate towards a decrease was more evident under 1 µM. This strategy allowed us to repeat twice the photocontrol of cortical activity by subsequent application of 1 µM *cis*-PAI and photoconversion to *trans*-PAI. The extent to which the diffusion of the drug reaches in adequate concentrations the deep layers of the cortex and to which the light reaches such deep layers needs further studies, given that the “engine” of slow oscillations lies in deep, layer 5-layer 6 layers.^42,47–50^

**Figure 4:**
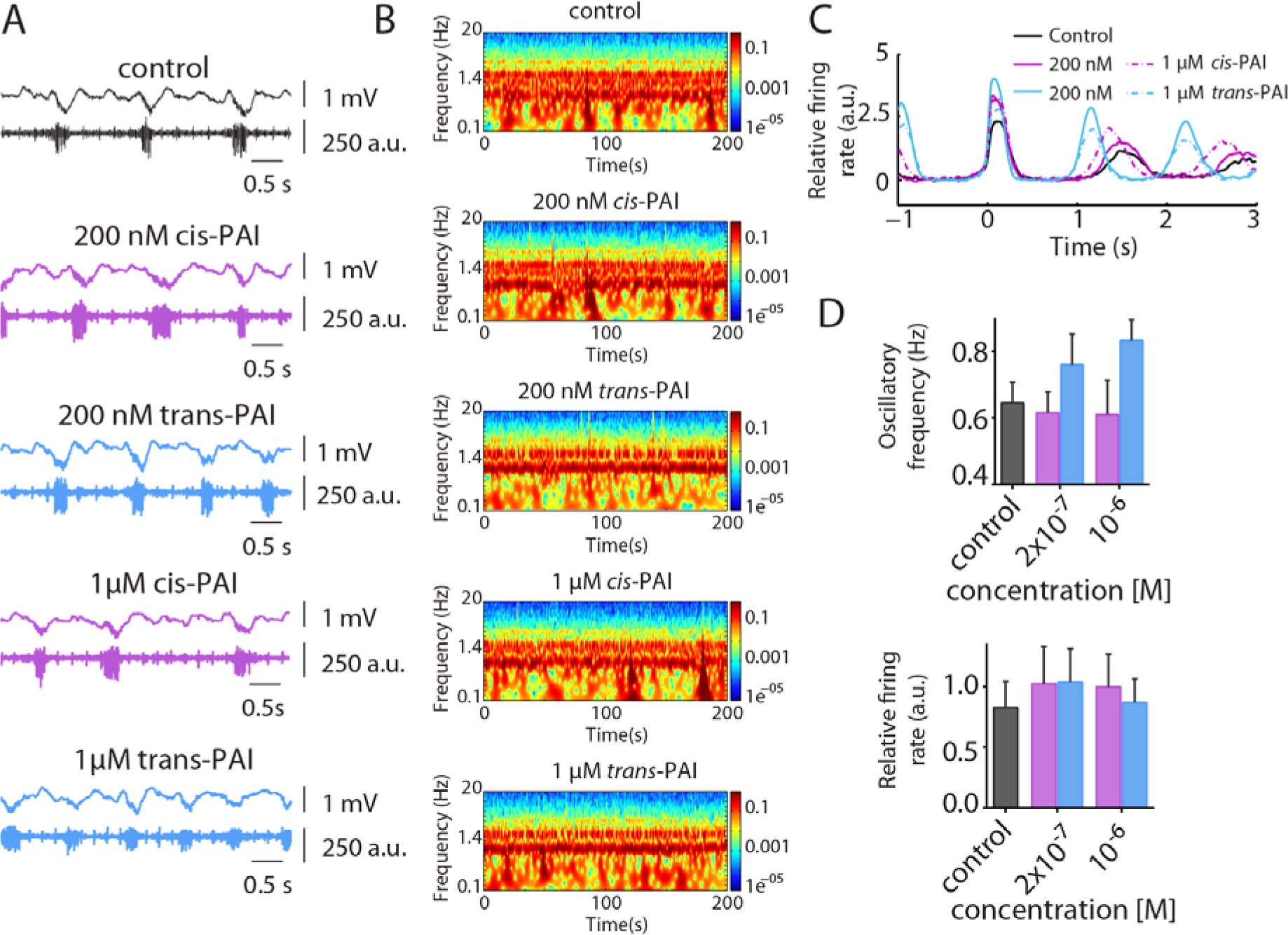
*In vivo* photomodulation of brain waves. **(A)** Representative raw traces of LFP (top) and Multiunit Activity (bottom), showing the differences in OF and relative firing rate between the control, 200 nM and 1 µM *cis*-PAI (pre-irradiated at 365 nm) and photoswitching with white light (*trans*-PAI). **(B)** Representative spectrogram under control, 200 nM and 1 µM *cis*-PAI (pre-irradiated at 365 nm) and photoswitching with white light (*trans*-PAI). **(C)** Representative examples of the waveform average of the MUA signal aligned at the Down-to-Up state transition under control, *cis*-PAI and *trans*-PAI at different concentrations (see legend). **(D)** Quantification of OF (Hz) and relative firing rate (Hz) at different concentrations.

## DISCUSSION

Different brain states are associated with distinct behaviors. In order to investigate the causality between them, behavioral outcomes must be correlated with the spontaneous and evoked activity in the cortical network. Thus, understanding the mechanisms of brain and behavioral state transitions requires new techniques for the manipulation of neuronal activity.^1^ They must enable the activation and inhibition of specific brain areas and neuronal circuits defined by several complementary criteria, namely electrical stimulation in selected regions, and photomanipulation that is cell-specific, or neurotransmitter-specific (using optogeneticc and photopharmacology, respectively).

Modulation of brain waves is also therapeutically useful in the treatment of CNS diseases, as shown by pioneering techniques used to control cortical activity noninvasively based on electromagnetic stimulation.^51,52^ Transcranial current stimulation enhances motor learning capacity in humans by cross-modulating the oscillatory neural activity (alpha and beta frequencies) in the motor cortex.^53–57^ Transcranial magnetic stimulation can normalize excessive gamma oscillations in the prefrontal cortex, and restore cognitive performance in patients with schizophrenia, Alzheimer’s and Parkinson’s diseases,^58,59^ and anxiety disorders.^60,61^ However, the physiological mechanisms of brain wave modulation are not known, and they are essential if we are to improve its spatiotemporal and spectral performance, both for fundamental and therapeutic purposes.^62^

Optogenetics^63–66^ has emerged as an alternative to controlling brain waves with electromagnetic fields: photocontrolling the release of ACh, which strongly modulates the transitions between different brain states,^9,12^ is possible by overexpressing photosensitive proteins in cholinergic neurons of mice neocortex.^10,67,68^ However, genetic manipulation is required in this approach. The light-dependent cholinergic muscarinic ligand presented here is, to date, the only way to directly photomodulate cholinergic pathways in intact tissue. We first studied the effect of the superagonist Iperoxo^40^ on isolated cortical slices (Fig. 1) in order to demonstrate that deep-sleep brain states can be controlled by selectively manipulating muscarinic receptors at their physiological location and context. The oscillatory frequency of the network was already greatly increased at 100 nM Iperoxo, and led to periods of seizure-like discharges, in agreement with previous studies performed using knockout mice and pilocarpine^44,45^ in which activation of muscarinic acetylcholine receptors induced seizures. Photocontrol of muscarinic signaling was subsequently achieved *in vitro* and *in vivo* with the photochromic iperoxo derivative PAI,^26^ which is targeted allosterically at M2 subtype receptors (Fig. 2–4). In particular, PAI is a dualsteric ligand displaying full agonism, which is probably important to achieve effective photocontrol in slices, where axons from cholinergic afferents have been cut and the concentration of ACh can be assumed to be zero due to the degradation by extracellular esterases. M2 mAChRs are involved in several CNS diseases like major depressive^25,69^ and bipolar disorders,^25,70^ Parkinson’s^25,71,72^ and Alzheimer’s^25,72,73^ diseases, but also in alcohol, smoking and drug dependence.^25,74,75^ These disorders are thus susceptible to drug-based photomodulation *in vivo* without requiring genetic manipulation. Although PAI cannot cross the blood-brain barrier due to its charge, and its safety profile has not been systematically characterized, it is less likely to trigger immune reactions and mutagenesis than microbial opsins.

In summary, the manipulation of brain state transitions, by means of photocontrolling the frequency of cortical oscillations, has been achieved with a photoswitchable dualsteric agonist of M2 mAChRs. This result opens the way to (1) dissecting the spatiotemporal distribution and pharmacology of brain states, namely how they depend on agonists, antagonists, and modulators of the different muscarinic subtypes expressed in the CNS, and (2) investigating the neuronal dynamics that regulate brain state transitions in the cortical surface and beyond. In particular, two-photon stimulation of PAI using pulsed infrared light^26^ offers the promise of subcellular resolution in three dimensions,^76^ as recently demonstrated with endogenous mGlu. ^77^ Compared to the local and often inhomogeneous expression patterns achieved with viral injections of optogenetic constructs, diffusible small molecules like PAI can in principle be applied to larger brain regions to control neuronal oscillations.^68^ Thus, remote control of brain waves based on the photopharmacological manipulation of endogenous muscarinic receptors may reveal the complex 3D molecular signaling underlying brain states and their transitions, in order to link them with cognition and behavior in a diversity of wild-type organisms.

## EXPERIMENTAL PROCEDURES

### Slice Preparation

Twenty-five ferrets (4- to 6-month-old) were anesthetized with sodium pentobarbital (40□mg/kg) and decapitated. The entire forebrain was rapidly removed and placed in oxygenated cold (4–10□°C) bathing medium.^78^ Ferrets were treated in accordance with protocols approved by the Animal Ethics Committee of the University of Barcelona, which comply with the European Union guidelines on the protection of vertebrates used for experimentation (Directive 2010/63/EU of the European Parliament and the Council of 22 September 2010). Coronal slices (400 µm thick) from primary visual cortex (V1) were used.^79^ To increase tissue viability we used a modification of the sucrose-substitution technique.^80^ During slice preparation, the tissue was placed in a solution in which NaCl was replaced with sucrose while maintaining the same osmolarity. After preparation, the slices were placed in an interface-style recording chamber (Fine Sciences Tools, Foster City, CA, USA). During the first 30□min the cortical slices were superfused with an equal mixture in volume of the normal bathing medium, artificial cerebral spinal fluid (ACSF) and the sucrose-substituted solution. Following this, normal bathing medium was added to the recording chamber and the slices were superfused for 1–2□h; the normal bathing medium contained (in mM): NaCl, 126; KCl, 2.5; MgSO_4_, 2; Na_2_HPO_4_, 1; CaCl_2_, 2; NaHCO_3_, 26; dextrose, 10; and was aerated with 95% O_2_, 5% CO_2_ to a final pH of 7.4. Then, a modified slice solution was used throughout the rest of the experiment; it had the same ionic composition except for different levels of the following (in mM): KCl, 4; MgSO_4_,1; and CaCl,1.^78^ Bath temperature was maintained at 34–36□°C.

### Drug application and photostimulation in brain slices

Iperoxo and PAI, both prepared as previously reported from commercially available starting materials,^26^ were bath-applied at the concentrations range of 1 nM to 100 nM for Iperoxo and 10 nM to 1 µM for PAI, as is mentioned in the Results section. We typically waited more than 1000 s after the application of the drug in order to let it act, to obtain a stable pattern of electrical activity, and to ensure a stable concentration in the bath. PAI effectively photomodulates the activity of M2 receptors *in vitro* and *in vivo*: in its dark-adapted state (*trans* form) it behaves as a strong M2 agonist, then upon illumination with UV light (365 nm), PAI switches to its off-state (*cis* form). PAI can be switched back to its on-state with blue or white light, or using two-photon excitation with pulsed NIR light.^26^ The high thermal stability of PAI inactive form (*cis* isomer) allows the administration of the inactive drug and subsequent activation of M2 receptors in the target region with white light.^26^ We first investigated the efficacy of PAI in cortical neuronal circuits *in vitro* by obtaining the dose-response curves of *trans*- and *cis*-PAI solutions applied separately. The more active PAI isomer (*trans*) was tested by applying its dark-adapted form (87 % of *trans*-PAI), and *cis*-PAI was obtained by illuminating 1 mM stock solutions with 365 nm light for 10 min (73% of *cis*-PAI).^26^ Increasing concentrations of both *trans*- and *cis*-PAI (10 nM, 100 nM, 300 nM and 1 µM) were bath applied in order to build up the dose-response curves.

### LFP recording and data analysis from *in vitro* recordings

LFP recordings started after allowing at least 2 h of recovery of the slices. Extracellular multiple unit recordings were obtained with flexible arrays of 16 electrodes arranged in columns as in Figure 1A. The multielectrode array (MEA) covered most of the area occupied by a cortical slice. It consisted of six groups of electrodes positioned to record electrophysiological activity from superficial and from deep cortical layers (692 μm apart) and from what should correspond to three different cortical columns (1500 μm apart). The unfiltered field potential (raw signal) was acquired at 10 kHz with a Multichannel System Amplifier (MCS, Reutlingen, Germany) and digitized with a 1401 CED acquisition board and Spike2 software (Cambridge Electronic Design, Cambridge, UK). The multiunit activity (MUA) was estimated as the power change in the Fourier components at high frequencies from the recorded LFP.^81^ Up-state detection was performed by setting a threshold in the log(MUA) time series as previously described to quantify frequency of the slow oscillations.^6,43^ Relative firing rate (FR) of the Up-states were quantified from the transformed log(MUA) signal as mean of absolute value of log(MUA). To study the variability of power spectral densities (PSD) of the local field potential, we used Welch’s method with 50% overlapped Hamming window with a resolution of 1 Hz. All off-line estimates and analyses were implemented in MATLAB (The MathWorks Inc., Natick, MA, USA). All variables in the experimental conditions were compared with the control (no chemical added) condition.

### The *in vivo* preparation

Cortical electrophysiology experiments were carried out in 2-3-month-old C57BL6/JR mice (n = 4) in accordance with the European Union Directive 2010/63/EU and approved by the local ethics committee. Mice were kept under standard conditions (room temperature, 12:12-h light-dark cycle, lights on at 08:00 a.m). Anesthesia was induced by intraperitoneal injection of ketamine (30 mg/kg) and medetomidine (100 mg/kg). After this procedure, the mouse was placed in a stereotaxic frame, and air was enriched with oxygen. Body temperature was maintained at 37°C throughout the experiment.^6^ A craniotomy was performed in each mouse: AP −2.5 mm, L 1.5 mm (primary visual cortex, V1).^82^ Cortical recordings were obtained from infragranular layers with 1–2 MΩ single tungsten electrode insulated with a plastic coating except for the tip (FHC, Bowdoin, ME, USA). Spontaneous local field potential (LFP) recordings from the V1 area provided information about the local neuronal population activity—within 250 μm.^83^ MUA estimation, Up-state detection and quantification of relative FR was performed as previously described. All these parameters were used to compare spontaneous activity during anesthesia (control), after application of the pre-illuminated, inactive drug form (*cis*-PAI) and application of white light to activate the drug (*trans*-PAI). *cis*-PAI was locally delivered to the cerebral cortex surface and activity was recorded while applying a commercial red filter on the white light source to avoid the activation of the drug. The uncovered brain was illuminated with a white light source (Photonic Optics™ Optics Cold Light Source LED F1) in order to activate the drug in situ (*trans*-PAI). The electrophysiological signal was amplified with a multichannel system (Multi Channel Systems), digitized at 20 kHz with a CED acquisition board and acquired with Spike 2 software (Cambridge Electronic Design) unfiltered.^84^

### Statistical analysis for slice experiments

Both *in vitro* and *in vivo* oscillatory frequency (OF) are reported as mean ± SEM. Measurements under different conditions were compared using the Friedman test and the Wilcoxon post-hoc tests corrected for multiple comparisons.^85^

## Acknowledgements

We thank Miquel Bosch for comments on the manuscript. This research received funding from European Union Research and Innovation Programme Horizon 2020 (Human Brain Project SGA2 Grant Agreement 785907, WaveScalES), European Research ERA-Net SynBio programme (Modulightor project), and financial support from Agency for Management of University and Research Grants/Generalitat de Catalunya (CERCA Programme; 2017-SGR-1442 project; RIS3CAT plan), Fonds Européen de Développement Économique et Régional (FEDER) funds, Ministry of Economy and Competitiveness (MINECO)/FEDER (Grant CTQ2016-80066-R to PG and BFU2017-85048-R to MVSV), and the Fundaluce foundation.

