## Supplementary Information for "Control of brain state transitions with light"

**Contents**

1. **Competition Binding experiments**
2. **Autocorrelograms**
1. **Competition Binding experiment**

**1.1 Materials and Methods**

Iperoxo (IPX) and Phthalimide-Azo-Iperoxo (PAI) were preliminarily assayed for their affinity to muscarinic receptors (mAChRs) by competition binding experiments in whole cortex of 3-4 months old female Wistar rat brain membrane, which contained a high density of all the subtypes of mAChRs.^1^ [^3^H]Quinuclidinyl benzilate ([^3^H]QNB) is a muscarinic antagonist without subtype selectivity, which binds the muscarinic receptors with high selectivity. [^3^H]QNB is recognized to be excellent for such binding experiments,^2,3^ and we performed them in order to test if IPX and PAI have the potential to modulate cortical brain states. Specific binding of IPX and PAI was defined by testing concentrations ranging from 10^-9^ to 10^-4^ M, and derivatizing the raw disintegrations per minute (dpm) data from the scintillation counter to obtain the total radioactivity values.^1^

**1.2 Results**

Competition binding experiments can show that both IPX and PAI have an interesting high binding affinity for mAChRs orthosteric site. IPX was found to have an IC_50_ of 1.5 µM, *trans*-PAI of 47 nM and *cis*-PAI of 118 nM (**Figure S1**). Such preliminary results encouraged us to investigate further our muscarinic compounds activity on the dynamics of the isolated V1 cortical slices.

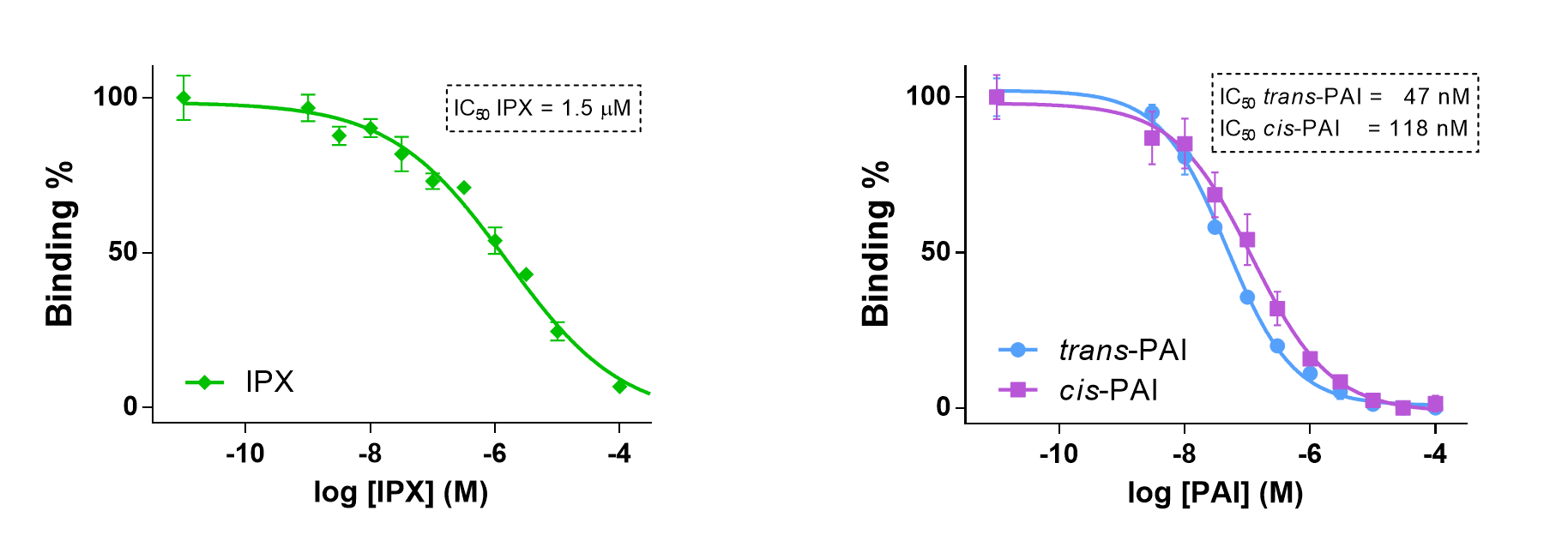

**Figure S1**. Competitive binding experiments of IPX and PAI to Wistar rats whole cortex containing all the mAChRs. Competition for specific binding of 200 pM [3H]QNB to 3-4 months-old female Wistar rats brain membranes (whole cortex) containing high density of all the five mAChRs by IPX and PAI (n = 4 for each isomer). Data points were fitted using the "log(inhibitor) vs. normalized response - Variable slope (four parameters" function in GraphPad Prism 6.

2. **Autocorrelograms**

**2.1 Autocorrelograms at 200 nM of PAI (switching of PAI activity)**

PAI (200 nM) was first applied in its inactive isomer (*cis*-PAI, pre-irradiation with 365 nm light). The activity of *cis*-PAI at 200 nM (pink line) in cortical slices does not evoke strong changes in terms of oscillatory activity in comparison to the basal control situation (black line) without PAI application, as it is shown in the autocorrelograms graph (**Figure S2**). After white light application, PAI switches to its active *trans* form (blue line), and strong changes of the oscillatory activity are visible (**Figure S2**). The autocorrelograms graphs are obtained by analyzing LFP from one channel.

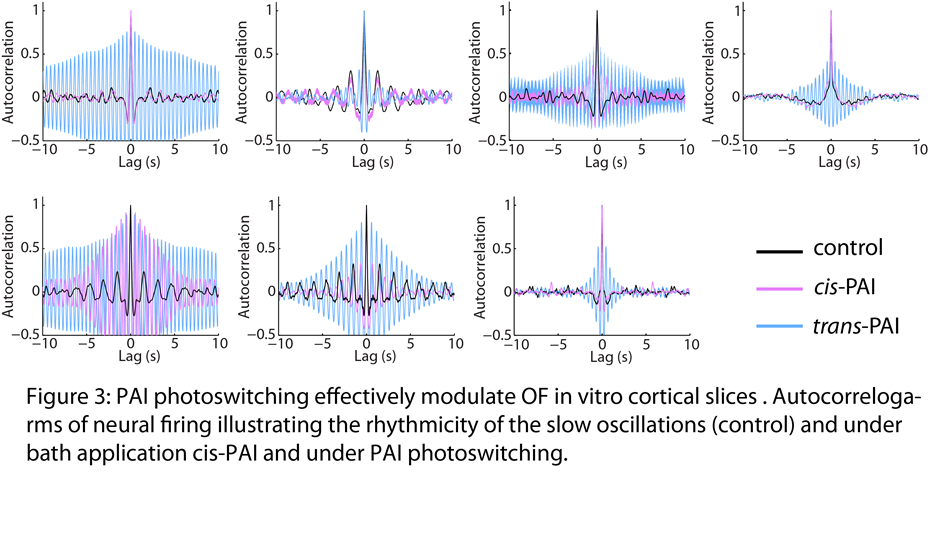

**Figure S2**. PAI can light-modulate the neuronal oscillatory activity in cortical ferret slices. Autocorrelograms of the rhythmicity of the neuronal oscillatory activity in basal condition (without PAI application, black line), under 200 nM *cis*-PAI application (pink line) and during white light irradiation (*trans*-PAI, blue line).

**2.2 Autocorrelograms at 100 nM and 1 µM of PAI**

The activity of PAI at 100 nM and 1 µM in cortical slices do not strongly differ between *trans* (**Figure S3**) and *cis* (**Figure S4**) in terms of oscillatory activity. 100 nM applications of *trans* and *cis* do not mostly produce changes in neuronal firing comparison to the basal control situation (black line), as it is shown in the autocorrelograms graph (**Figure S3** and **S4**). At 1 µM, both PAI isomers produce strong changes of the oscillatory activity (**Figure S3** and **S4**). The autocorrelograms graphs are obtained by analyzing LFP from one channel.

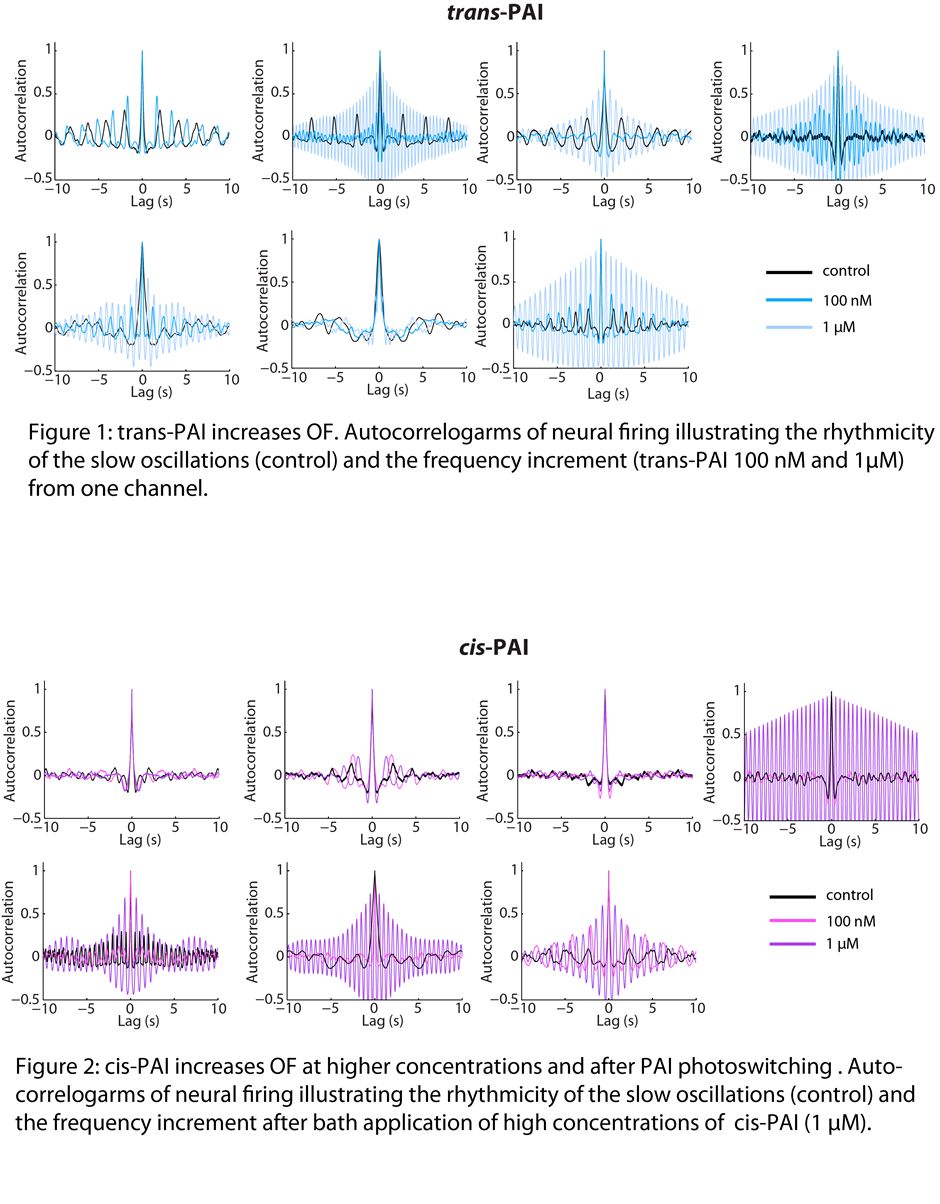

**Figure S3**. Autocorrelograms of the rhythmicity of the neuronal oscillatory activity in basal condition (without PAI application, black line), at 100 nM and 1 µM of *trans*-PAI (blue lines) applications.

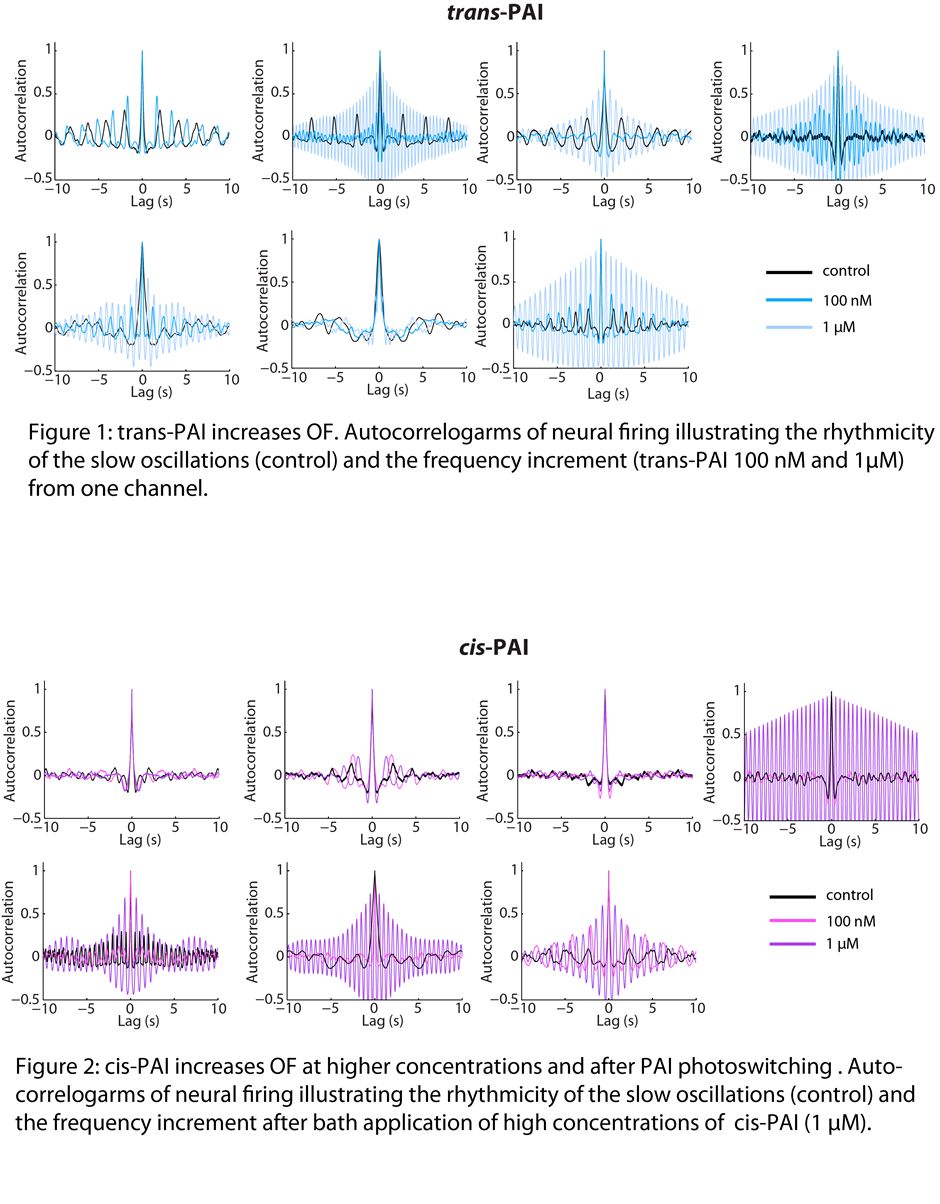

**Figure S4**. Autocorrelograms of the rhythmicity of the neuronal oscillatory activity in basal condition (without PAI application, black line), at 100 nM and 1 µM of *cis*-PAI (pink lines) applications.
